# Three models of vaccination strategies against cryptococcosis in immunocompromised hosts using heat-killed *Cryptococcus neoformans* Δ*sgl1*

**DOI:** 10.1101/2022.01.31.478598

**Authors:** Tyler G. Normile, Maurizio Del Poeta

## Abstract

Vaccines are one of the greatest medical accomplishments to date, yet no fungal vaccines are currently available in humans mainly because opportunistic mycoses generally occur during immunodeficiencies necessary for vaccine protection. In previous studies, a live, attenuated *Cryptococcus neoformans* Δ*sgl1* mutant accumulating sterylglucosides was found to be avirulent and protected mice from a subsequent lethal infection even in absence of CD4^+^ T cells, a condition most associated with cryptococcosis (e.g., HIV). Here, we tested three strategies of vaccination against cryptococcosis. First, in our preventative model, protection was achieved even after a 3-fold increase of the vaccination window. Second, because live *C. neoformans* Δ*sgl1*-vaccinated mice challenged more than once with WT strain had a significant decrease in lung fungal burden, we tested *C. neoformans* Δ*sgl1* as an immunotherapeutic. We found that therapeutic administrations of HK *C. neoformans* Δ*sgl1* subsequent to WT challenge significantly improve the lung fungal burden. Similarly, therapeutic administration of HK *C. neoformans* Δ*sgl1* post WT challenge resulted in 100% or 70% survival depending on the time of vaccine administration, suggesting that HK Δ*sgl1* is a robust immunotherapeutic option. Third, we investigated a novel model of vaccination in preventing reactivation from lung granuloma using *C. neoformans* Δ*gcs1*. Remarkably, we show that administration of HK Δ*sgl1* prevents mice from reactivating Δ*gcs1* upon inducing immunosuppression with corticosteroids or by depleting CD4^+^ T cells. Our results suggest that HK Δ*sgl1* represents a clinically relevant, efficacious vaccine that confers robust host protection in three models of vaccination against cryptococcosis even during CD4-deficiency.

**Importance:** Cryptococcosis results in ∼180,000 global deaths per year in immunocompromised individuals. Current antifungal treatment options are potentially toxic, lacking in areas of need, and exhibit limited efficacy. In addition to these lackluster therapeutic options, no fungal vaccines are currently available for clinical use. Due to the increasing rate of immunocompromised individuals, there is a dire need for the development of improved antifungal therapeutics. Presently, we have demonstrated the high efficacy of a clinically relevant heat-killed mutant strain of *Cryptococcus neoformans* in inducing advantageous host protection in three models of vaccination against cryptococcosis during immunodeficiencies most associated with this disease.

## Introduction

Invasive fungal infections are primarily caused by environmental fungi that mainly infect immunocompromised individuals resulting in ∼1.5 million deaths a year that account for ∼50% of all AIDS-related deaths [1, 2]. Individuals most at risk include HIV/AIDS patients [3–5], cancer patients receiving chemotherapy [6, 7], solid organ transplant recipients [8–10], or patients taking medication to control chronic diseases [11–14]. Unlike most fungi that do not infect humans, the pathogenicity of invasive fungal species begins with the ability to grow and replicate at human body temperature [15, 16], which suggests that climate change, particularly global warming, may play a role in increasing infections from environmental fungi in more temperate climates [17–19]. The incidence of invasive fungal infections is expected to further increase as the global immunocompromised population continues to rise due to novel immunosuppressive therapies or comorbidities, such as the current COVID-19 pandemic [20–25].

One of these fungal pathogens is *Cryptococcus neoformans*, a basidiomycetous yeast ubiquitously found in environmental sources such as avian habitation, trees, and soil [3, 5, 26]. *C. neoformans* is a main etiological agent of cryptococcosis, a life-threatening invasive fungal infection that originates in the respiratory tract [27–29]. Upon inhalation of the environmental fungal propagules, immunocompetent hosts often remain asymptomatic while they either clear the initial infection eliminating the yeast from the lungs or control fungal proliferation by enclosing the persistent yeast in lung granulomas where the fungal cells remain dormant [30–32]. Conversely, immunocompromised individuals lacking a necessary component of the immune system, namely CD4^+^ T cells as seen with HIV/AIDS, generally fail to control the initial infection or maintain the integrity of the lung granulomas containing latent cryptococcal cells leading to host pathology [11, 33]. These individuals may experience uncontrolled fungal replication and dissemination of the fungus to the central nervous system potentially leading to life-threatening meningoencephalitis [9, 33] accounting for ∼220,000 new cryptococcal cases and ∼180,000 deaths a year [34, 35].

Vaccines are considered to be one of the greatest medical accomplishments to date [36]. Although the high mortality rate upon extrapulmonary cryptococcosis in at-risk individuals can be partly attributed to the poor efficacy, host toxicity, and pathogen-acquired resistance of current antifungal therapeutics [37–39], the absence of fungal vaccines is a major constraint in overcoming invasive fungal infections in humans [40]. While there has been ample research into the development of a vaccine against *C. neoformans* (reviewed in [41–43]), none have advanced past the pre-clinical research stage. The lack of vaccine advancement is chiefly due to the fact this pathogen infects mostly immunocompromised individuals with low CD4^+^ T cell counts [3, 28], and the majority of current cryptococcal vaccine research lack host protective efficacy in this immunodeficiency. As such, vaccine formulations exhibiting high efficacy in animal models that resemble immunodeficiencies associated with cryptococcosis (e.g., lacking CD4^+^ T cells) are in high demand [42, 44].

Exposure to *C. neoformans* may result in the yeast being cleared or safely contained within lung granulomas in immunocompetent hosts [31, 33]. In addition to the necessity of vaccine studies being carried out in immunodeficient conditions, the literature currently contains only cryptococcal vaccines used in a prophylactic manner. However, there are no reports of vaccination strategies against the reactivation of dormant *C. neoformans* from lung granuloma breakdown due to immunosuppressive occurrences (reviewed in [44]). This disparity is mainly attributed to the lack of tools to evaluate vaccines against infection by the reactivation of latent fungal cells in mouse models since mice do not form lung granulomas to wild-type (WT) *C. neoformans* and remains a major understudied bottleneck in the advancement of a clinically available anti-cryptococcal vaccine.

Our lab has previously engineered a mutant strain of *C. neoformans* (Δ*sgl1*) that accumulates large amounts of sterylglucosides (SGs) and provided the first evidence on the key role of sterylglucosidase 1 (Sgl1) on fungal virulence [45]. SGs have been previously shown to possess immunological functions (reviewed in [46]). The use of the plant SG, β-sitosterol, increased T cell proliferation and Th1 polarization [47, 48], significantly prolonged survival of mice infected with *Candida albicans* [49, 50], and promoted the recovery of patients with pulmonary tuberculosis in combination with regular anti-tuberculosis treatment [48]. However, our work provided the first physio-pathological studies with fungal-derived SGs, and our recent structural studies will enable the rational design of new antifungal agents targeting Sgl1 [51].

Prior studies in our lab have shown that *C. neoformans* Δ*sgl1* induces a proinflammatory lung cytokine environment with robust effector cell recruitment to the lungs as well as confers complete host protection to lethal WT challenge under immunodeficiencies most associated with cryptococcosis (e.g., lacking CD4^+^ T cells) [52]. Interestingly, we found that protection required SGs in combination to the immunosuppressive glucuronoxylomannan (GXM)-based capsule since an acapsular mutant strain (Δ*cap59*Δ*sgl1*) was no longer protective [53] nor induce aprotective cytokine response to *ex vivo* stimulated γ*δ* T cells (T.G. Normile, T.H. Chu, B.S. Sheridan, and M. Del Poeta, submitted for publication), suggesting that SGs may act as an immunoadjuvant to GXM to induce host protection. Most recently, we uncovered the immune-mechanism of protection involved TLR2-mediated production of IFNγ and IL-17A by γ*δ* T cells resulting in a robust CD4^+^ or CD8^+^ T cell response for complete host protection to subsequent WT infection (T.G. Normile, T.H. Chu, B.S. Sheridan, and M. Del Poeta, submitted for publication). Overall, these studies suggest that *C. neoformans* Δ*sgl1* represents a viable live, attenuated vaccine.

In the present study, we validate three different models of successful vaccination strategies against cryptococcosis using heat-killed (HK) *C. neoformans* Δ*sgl1* in condition of CD4^+^ T cell deficiency. In the canonical prevention model of vaccination, we found that two subsequent administrations of HK *C. neoformans* Δ*sgl1* conferred complete host protection to a WT challenge even when CD4^+^ T cells were depleted, mimicking the results obtained with the live, attenuated mutant. Host protection in immunocompetent and CD4-deficient mice was still found after increasing the time of the vaccination window from 30 to 90 days or after challenging vaccinated mice 3 subsequent times, suggesting our vaccine strategy induces a long-lasting and non-exhaustive memory T cell protection. Interestingly, vaccinated mice receiving multiple WT challenges showed a significant decrease in lung fungal burden compared to vaccinated mice that were challenged only once. Because of these findings, we tested whether *C. neoformans* Δ*sgl1* could be used in a therapeutic manner. We found that administration of HK *C. neoformans* Δ*sgl1* post WT challenge in naive mice significantly prolonged survival compared to untreated mice. In previously vaccinated mice, administration of HK *C. neoformans* Δ*sgl1* post WT challenge significantly decreased the lung fungal burden post challenge, even during CD4^+^ T cell deficiency. Finally, we tested HK *C. neoformans* Δ*sgl1* in a model of cryptococcal granuloma to study whether our vaccination strategy would prevent fungal reactivation upon immunosuppression. We found that *C. neoformans* Δ*sgl1*-vaccinated mice exhibited significantly enhanced survival and control of fungal proliferation from latent granuloma-contained fungal cells upon inducing immunosuppression with either corticosteroid administration or CD4^+^ T cell depletion.

In conclusion, our results suggest that HK *C. neoformans* Δ*sgl1* represents a clinically relevant vaccine candidate and confers robust host protection in three models of vaccination against cryptococcosis during host conditions most associated with clinical cases of cryptococcosis in humans.

## Results

### Vaccination with HK C. neoformans Δsgl1 confers concentration-dependent host protection

We have recently found that murine splenocytes robustly produced the essential protective cytokines IFNγ and IL-17A when stimulated with HK *C. neoformans* Δ*sgl1 ex vivo*, and in fact *ex vivo* stimulation of splenocytes produced significantly greater quantities of these cytokines compared to live-cell stimulation at the same concentration and on the same timeline (T.G. Normile, T.H. Chu, B.S. Sheridan, and M. Del Poeta, submitted for publication). From this observation, we asked if administration of HK *C. neoformans* Δ*sgl1* would provide the same host protection to lethal WT challenge as with the live, attenuated mutant.

Since vaccination with HK mutant strains is notoriously known to elicit a weaker immune response than live, attenuated strains [54], mice were administered increasing concentrations of HK *C. neoformans* Δ*sgl1* 30 days prior to WT challenge. As expected, mice vaccinated with live *C. neoformans* Δ*sgl1* were fully protected while unvaccinated mice fully succumbed to infection (**Fig. 1A**). Interestingly, we observed a concentration-dependent survival rate in mice with the increasing concentrations of HK *C. neoformans* Δ*sgl1*. Mice administered 5x10^5^ HK *C. neoformans* Δ*sgl1* fully succumbed to the WT infection in a similar timeline to unvaccinated mice (**Fig. 1A**). There was a significant increase in median survival time for mice administered 5x10^6^ HK *C. neoformans* Δ*sgl1*, although all mice still succumbed to infection. Remarkably, mice administered 5x10^7^ HK *C. neoformans* Δ*sgl1* exhibited a 70% survival rate at the endpoint of the experiment that was not statistically different from the complete protection seen with live *C. neoformans* Δ*sgl1* (**Fig. 1A**).

**Figure 1.**
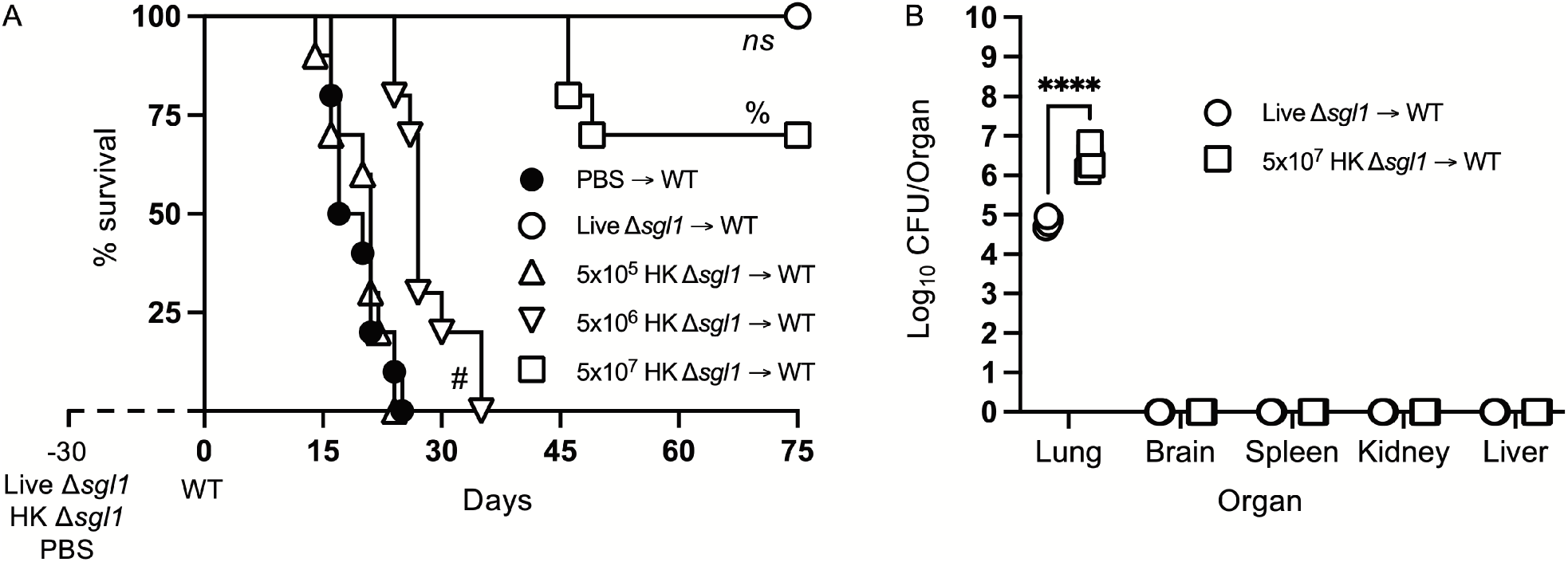
Vaccination with heat-killed (HK) *C. neoformans* Δ*sgl1* confers concentration-dependent partial host protection. **A.** CBA/J mice (n=10 mice/group) were administered 5x10^5^ Live Δ*sgl1*, 5x10^5^, 5x10^6^, or 5x10^7^ HK Δ*sgl1*, or PBS. After 30 days, mice were challenged with 5x10^5^ *C. neoformans* wild-type (WT) (day 0) and monitored for survival. **B.** Endpoint organ fungal burden was quantified in the lungs, brain, spleen, kidney, and liver in surviving mice (n=4 mice/group). Graphed data represent the survival percentage of WT challenged mice (**A**) or the mean +/-SD (**B**). Significance was determined using a two-tailed unpaired t-test (**B**) and significance is denoted as ****, *P* < 0.001. Survival significance was determined by the Mantel-Cox log-rank test (**A**) and denoted on graph **A**: *ns*, not significant (*P* > 0.05) for Live Δ*sgl1* → WT vs. 5x10^7^ HK Δ*sgl1* → WT; %, *P* < 0.001 for 5x10^7^ HK Δ*sgl1* → WT vs. 5x10^6^ HK Δ*sgl1* → WT; #, *P* < 0.001 for 5x10^6^ HK Δ*sgl1* → WT vs. 5x10^5^ HK Δ*sgl1* → WT.

Surviving mice visually appeared healthy with normal weight gain, and endpoint organ fungal burden analysis confirmed no extrapulmonary dissemination had occurred (**Fig. 1B**). Nevertheless, mice vaccinated with 5x10^7^ HK *C. neoformans* Δ*sgl1* displayed a significantly greater lung fungal burden compared to mice vaccinated with live *C. neoformans* Δ*sgl1* (**Fig. 1B**). These data suggest that mice vaccinated with HK *C. neoformans* Δ*sgl1* exhibited concentration-dependent partial protection with 5x10^7^ HK *C. neoformans* Δ*sgl1* being the most efficacious.

### Two administrations of HK C. neoformans Δsgl1 confers complete host protection even during CD4^+^ T cell immunodeficiency

We have unveiled that administration of a single dose of 5x10^7^ HK *C. neoformans* Δ*sgl1* conferred similar host protection compared to vaccination with live *C. neoformans* Δ*sgl1*, although complete protection was not achieved (**Fig. 1A**), and the endpoint lung fungal burden was significantly greater than the live mutant vaccinated mice (**Fig. 1B**). However, due to the decreased length of antigen encounter, vaccination with HK mutants offer less host cell stimulation of protective cytokines, decreased naïve T cell expansion, and attenuated memory T cell formation [36, 54].

Since we have previously reported that either CD4^+^ or CD8^+^ T cells are required for *C. neoformans* Δ*sgl1*-mediated host protection [52], we hypothesized that repeated immunization with this HK mutant dose may negate the negative facets of HK vaccination and promote stronger adaptive T cell-mediated immunity as seen with other HK mutant vaccine studies [55, 56]. We tested this hypothesis by administering two subsequent doses of 5x10^7^ HK *C*. *neoformans* Δ*sgl1* (HK2d Δ*sgl1*) on days -30 and -15 prior to the WT challenge. Indeed, mice that received two administrations of 5x10^7^ HK *C. neoformans* Δ*sgl1* exhibited complete host protection (100% survival) at the experimental endpoint (**Fig. 1A**). Endpoint organ fungal burden analysis showed that HK2d Δ*sgl1*-vaccinated mice displayed no extrapulmonary dissemination and a significantly lower lung fungal compared to live *C. neoformans* Δ*sgl1*-vaccinated mice (**Fig. 1B**). In fact, 1 of the 7 HK2d Δ*sgl1*-vaccinated mice fully cleared the WT yeast from the lungs. These data suggest that vaccination with two subsequent doses of 5x10^7^ HK *C. neoformans* Δ*sgl1* confers complete host protection and aids in pulmonary clearance of the WT fungal cells.

**Figure 2.**
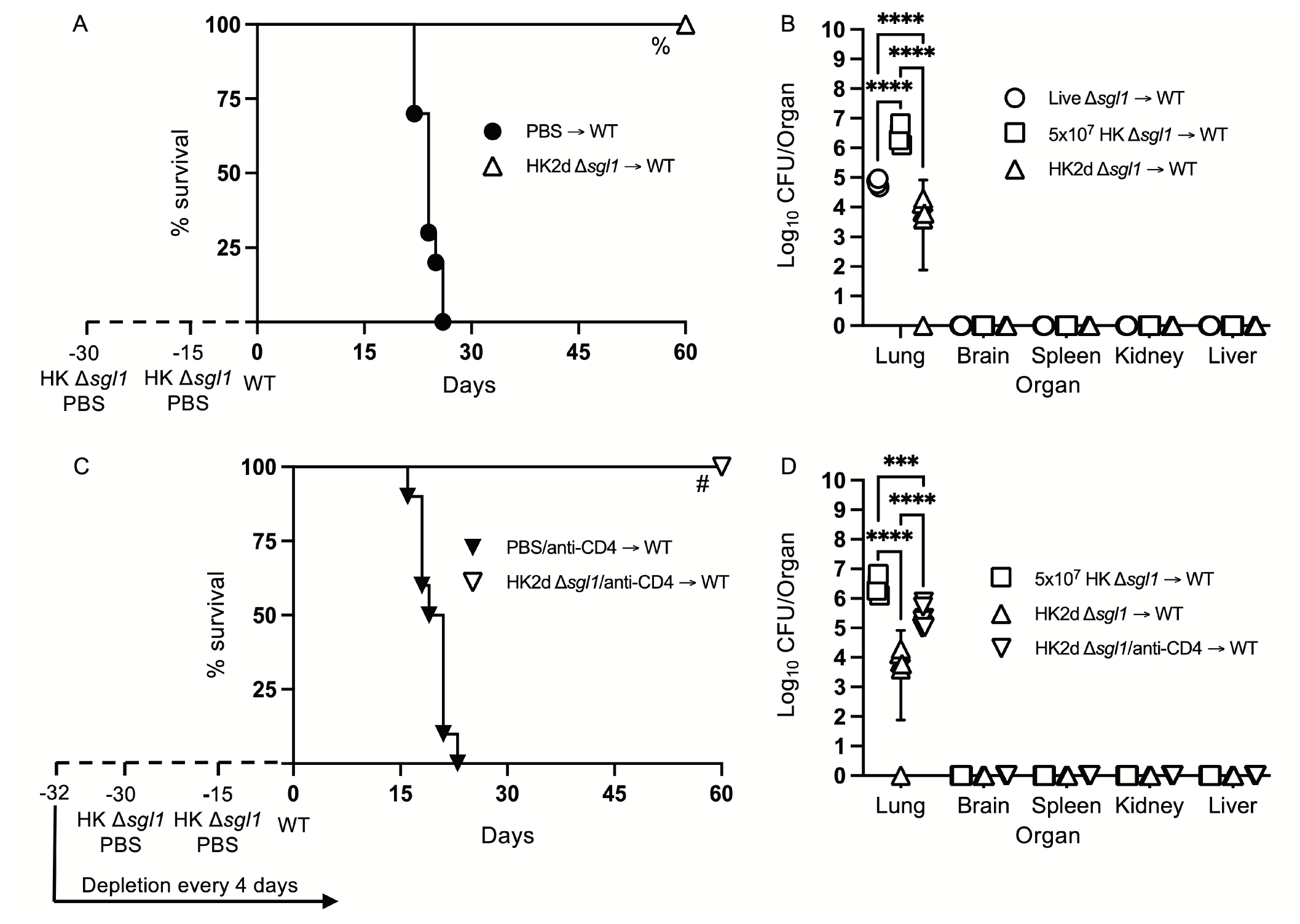
Two doses of heat-killed (HK2d) *C. neoformans* Δ*sgl1* confers complete host protection even in the absence of CD4^+^ T cells. **A.** CBA/J mice (n=10 mice/group) were administered two identical doses of 5x10^7^ HK Δ*sgl1* or PBS (days -30 and -15), challenged with 5x10^5^ *C. neoformans* wild-type (WT) (day 0), and monitored for survival. **B.** Endpoint organ fungal burden was quantified in the lungs, brain, spleen, kidney, and liver from HK2d Δ*sgl1* → WT and compared to the endpoint fungal burden in Live Δ*sgl1* → WT and 5x10^7^ HK Δ*sgl1* → WT from Figure 1B (n=4-7 mice/group). **C.** CBA/J mice (n=10 mice/group) were depleted of CD4^+^ T cells prior to administration of two identical doses of 5x10^7^ HK Δ*sgl1* or PBS (days -30 and -15), challenged with 5x10^5^ *C. neoformans* WT (day 0), and monitored for survival. **D.** Endpoint organ fungal burden was quantified in the lungs, brain, spleen, kidney, and liver in HK2d Δ*sgl1*/anti-CD4 → WT mice and compared to the endpoint fungal burden from 5x10^7^ HK Δ*sgl1* → WT and HK2d Δ*sgl1* → WT from Figure 2B (n=4-7 mice/group). Graphed data represent the survival percentage of WT challenged mice (**A, C**) or the mean +/-SD (**B, D**). Significance was determined by a two-way ANOVA using Šídák’s multiple comparisons test for *P* value adjustment (**B, D**) and significance is denoted as ***, *P* < 0.005; ****, *P* < 0.001. Survival significance was determined by the Mantel-Cox log-rank test (**A, C**) and denoted on each graph: **A**: %, *P* < 0.001 for HK2d Δ*sgl1* → WT vs. PBS → WT; **C**: #, *P* < 0.001 for HK2d Δ*sgl1*/anti-CD4 → WT vs. PBS/anti-CD4 → WT.

To assess if vaccination with HK *C. neoformans* Δ*sgl1* possessed clinical relevance, CD4-deficient mice were also vaccinated with two subsequent doses of 5x10^7^ HK *C. neoformans* Δ*sgl1* and challenged mice with the WT strain. Interestingly, 100% host protection was achieved in HK2d Δ*sgl1*-vaccinated mice depleted of CD4^+^ T cells (**Fig. 1C**), and endpoint organ fungal burden analysis revealed no extrapulmonary dissemination of the WT strain (**Fig. 1D**). There was a significantly greater fungal burden in the lungs of HK2d Δ*sgl1*-vaccinated mice depleted of CD4^+^ T cells compared to HK2d Δ*sgl1*-vaccinated immunocompetent mice. However, there was a significantly lower lung fungal burden in HK2d Δ*sgl1*-vaccinated mice depleted of CD4^+^ T cells compared to live *C. neoformans* Δ*sgl1*-vaccinated immunocompetent mice (**Fig. 1D**).

Overall, these data indicate that vaccination with two subsequent administrations of 5x10^7^ HK *C. neoformans* Δ*sgl1* confers host protection from WT challenge in both immunocompetent and CD4-deficient mice, and the HK vaccination strategy may provide a greater efficacy in host clearance of the WT strain from the lungs compared to live vaccination strategy.

### Vaccination with live or HK C. neoformans Δsgl1 confers long-lasting host immunity to lethal WT infection

Because administration of 2 subsequent doses of 5x10^7^ HK *C. neoformans* Δ*sgl1* also conferred complete host protection to the WT strain even during CD4-deficiency (**Fig. 1**), we sought to investigate the efficacy of host protection after vaccination with either live or HK *C. neoformans* Δ*sgl1* via alternations to our preventative vaccination model during immunocompetency and CD4-deficiency.

To assess the longevity of the vaccine-induced memory T cells, we increased the time between the administration of the vaccine and WT challenge. Immunocompetent or CD4-deficient mice were administered either live *C. neoformans* Δ*sgl1* or PBS and challenged with the WT strain 90 days later (a 3-fold increase between vaccination and WT challenge). Interestingly, all vaccinated mice survived the WT challenge, while all unvaccinated mice succumbed to the WT infection (**Fig. 2A**). Endpoint organ fungal burden in surviving mice showed that no extrapulmonary dissemination was observed (**Fig. 2B**). Similar to our previous studies, the lung fungal burden in CD4-deficient vaccinated mice was significantly greater than in immunocompetent vaccinated mice. These data show that vaccination with live *C. neoformans* Δ*sgl1* confers long term host immunity to lethal WT challenge, which strongly suggests long-lived memory T cells even during CD4-deficiency.

**Figure 3.**
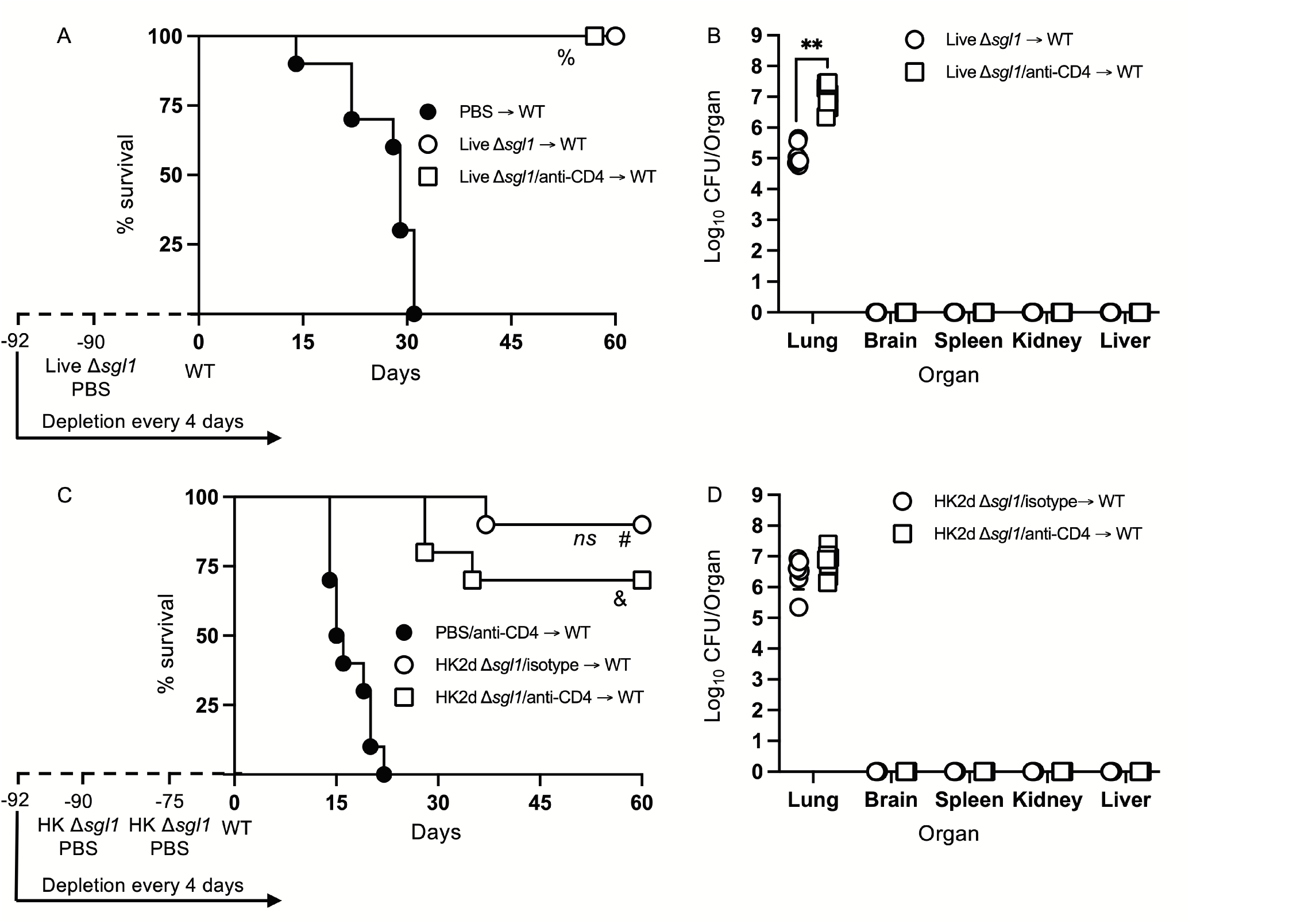
Vaccination with live or heat-killed (HK) *C. neoformans* Δ*sgl1* confers long-lasting host protection. **A.** CBA/J mice (n=10 mice/group) were administered anti-CD4 antibody or left untreated prior to vaccination with 5x10^5^ Live Δ*sgl1* or PBS controls, and the depletions continued for the entirety of the experiment at noted intervals. Mice were given an extended 90-day rest period where vaccinated and unvaccinated mice were then challenged with 5x10^6^ *C. neoformans* wild-type (WT) (day 0) and monitored for survival. **B.** Endpoint organ fungal burden was quantified in the lungs, brain, spleen, kidney, and liver in surviving mice (n=8 mice/group). **C.** CBA/J mice (n=10 mice/group) were administered either isotype or anti-CD4 antibodies prior to vaccination with two identical doses of 5x10^7^ HK Δ*sgl1* or PBS controls on days -90 and -75, and the depletions continued for the entirety of the experiment at noted intervals. Mice were given an extended 90-day rest period where vaccinated and unvaccinated mice were then challenged with 5x10^6^ *C. neoformans* WT (day 0) and monitored for survival. **D.** Endpoint organ fungal burden was quantified in the lungs, brain, spleen, kidney, and liver in surviving mice (n=7 mice/group). Graphed data represent the survival percentage of WT challenged mice (**A, C**) or the mean +/- SD (**B, D**). Significance was determined by a two-tailed unpaired t-test (**B, D**) and significance is denoted as **, *P* < 0.01. Survival significance was determined by the Mantel-Cox log-rank test (**A, C**) and denoted on each graph: **A**: %, *P* < 0.001 for Live Δ*sgl1* → WT or Live Δ*sgl1*/anti-CD4 → WT vs. PBS → WT; **C**: *ns*, not significant (*P* > 0.05) for HK2d Δ*sgl1*/isotype → WT vs. HK2d Δ*sgl1*/anti-CD4 → WT; #, *P* < 0.001 for HK2d Δ*sgl1*/isotype → WT vs. PBS/anti-CD4 → WT; &, *P <* 0.001 for HK2d Δ*sgl1*/anti-CD4 → WT vs. PBS/anti-CD4 → WT.

Because vaccination with live *C. neoformans* Δ*sgl1* promoted long term immunity resulting in complete host protection to the WT strain, we then asked if HK *C. neoformans* Δ*sgl1* provided the same protection. Immunocompetent or CD4-deficient mice were administered 2 subsequent doses of either HK *C. neoformans* Δ*sgl1* or PBS on days -90 and -75 and challenged with the WT strain on day 0. We observed a 90% and 70% survival rate in immunocompetent and CD4-deficient mice, respectively (**Fig. 2C**). Nonetheless, the difference between the median survival time for immunocompetent mice and CD4-deficient mice was not statistically different, the endpoint lung fungal burdens were nearly identical, and no extrapulmonary dissemination of the WT yeast was observed in either group (**Fig. 2D**). Altogether, these data suggest that vaccination with live or HK *C. neoformans* Δ*sgl1* provides long-lived memory T cells conferring robust host protection and lung containment even after a 3-fold increase of the vaccination window.

### Vaccination with C. neoformans Δsgl1 confers complete host protection to multiple WT challenges even in the absence of CD4^+^ T cells

Since vaccination with either live or HK *C. neoformans* Δ*sgl1* provides long-term immunity, we then wanted to investigate the possibility of induced T cell anergy. During chronic infections following an antigen encounter, T cells may become tolerant and non-responsive but remain alive for extended periods of time in a hyporesponsive state [57]. Due to the fact that we observe persistent fungal cells in the lungs post WT challenge, T cell anergy may potentially occur after the contraction phase post WT challenge.

To test for induced T cell anergy, immunocompetent or CD4-deficient *C. neoformans* Δ*sgl1*-vaccinated mice underwent multiple WT challenges, monitored for survival, and lung fungal burden was assessed at the end of each WT challenge period (experimental design schematic: **Fig. 3A**). Very interestingly, *C. neoformans* Δ*sgl1*-vaccinated mice completely survived for a total of 105 days after three subsequent lethal WT challenges on days 0, 45, and 75 (**Fig. 3B**). Endpoint lung fungal burden analysis showed that there was a significant decrease of persistent WT yeast in the lungs of mice that were WT challenged a second time (**Fig. 3C**). This decrease in lung burden did not further decrease after a third challenge. In addition, the decrease in the persistent lung fungal burden from the subsequent WT challenge resulted in no statistical difference between the lung burden in isotype-treated and CD4-deficient mice (**Fig. 3C**). Overall, these data indicate that immunocompetent and CD4-deficient *C. neoformans* Δ*sgl1*-vaccinated mice are protected from at least three subsequent lethal WT challenges resulting in fewer persisting WT cells in the lungs strongly suggesting that vaccination produces memory T cells that do not undergo anergy due to chronic infection.

**Figure 4.**
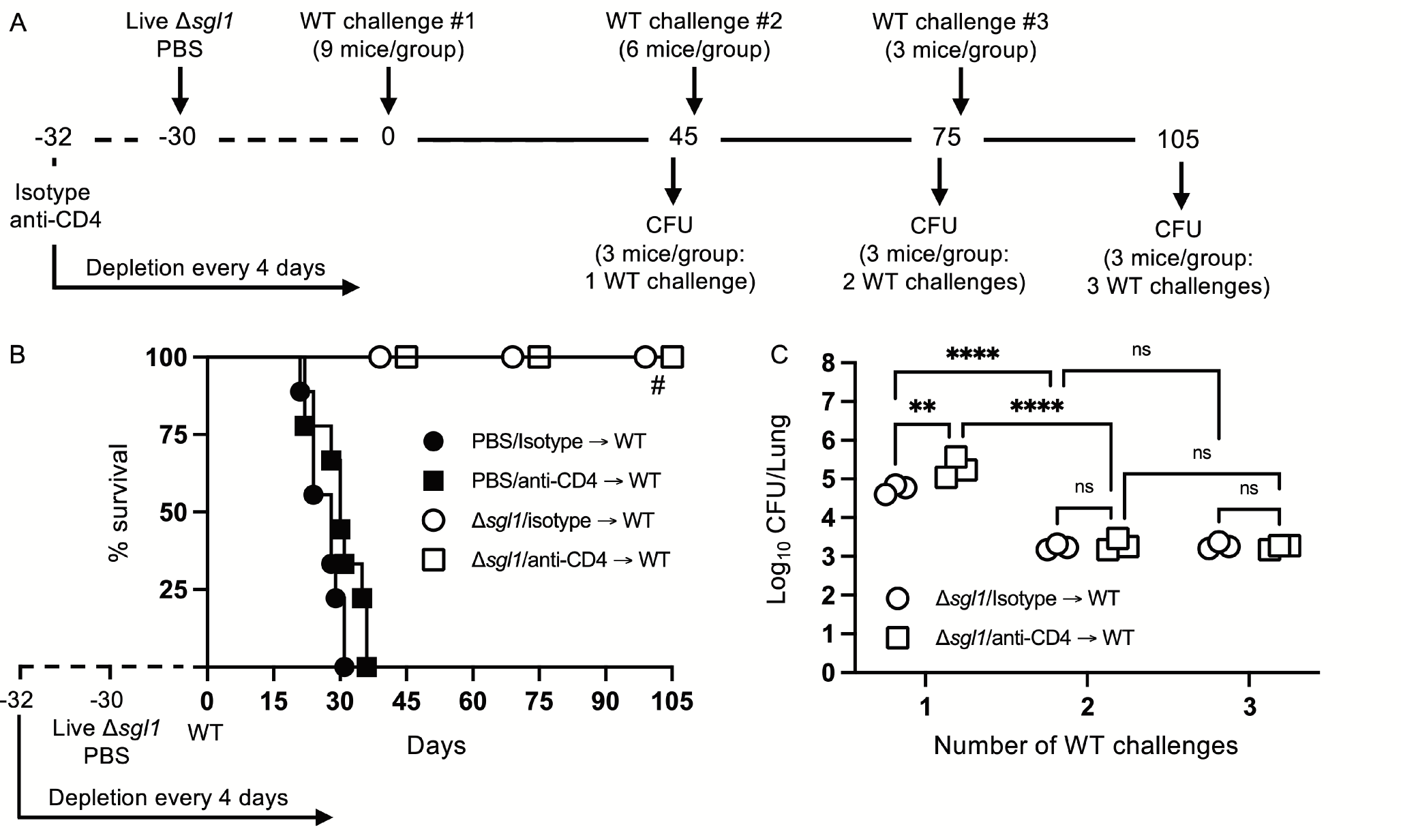
Vaccination with *C. neoformans* Δ*sgl1* confers complete host protection to multiple subsequent wild-type (WT) challenges even in the absence of CD4^+^ T cells. **A**. Experimental design schematic. **B.** CBA/J mice (n=9 mice/group) were administered either isotype or anti-CD4 antibody prior to vaccination with 5x10^5^ Live Δ*sgl1* or PBS controls, and the depletion continued for the entirety of the experiment at noted intervals. After 30 days, vaccinated and unvaccinated mice were challenged with 5x10^5^ *C. neoformans* WT on day 0 (WT challenge #1). Only vaccinated mice remained on day 45 post WT challenge #1, and 3 mice/group were sacrificed for organ fungal burden while the remaining mice were all challenged a second time with 5x10^5^ *C. neoformans* WT on day 45 (WT challenge #2). 30 days later on day 75, 3 mice/group were sacrificed for organ fungal burden while the remaining mice were all challenged a third time with 5x10^5^ *C. neoformans* WT on day 75 (WT challenge #3). Finally, 30 days later on day 105, the remaining 3 mice/group were sacrificed for organ fungal burden. **C.** Endpoint organ fungal burden was quantified in the lungs for mice sacrificed on days 45 (1 WT challenge), 75 (2 WT challenges), and 105 (3 WT challenges) (n=3 mice/group). Graphed data represent the survival percentage of WT challenged mice (B) or the mean +/- SD (**C**). Significance was determined by a two-way ANOVA using Šídák’s multiple comparisons test for *P* value adjustment (**C**) and significance is denoted as *ns*, not significant (*P* > 0.05); **, *P* < 0.01; ****, *P* < 0.001. Survival significance was determined by the Mantel-Cox log-rank test (**B**) and denoted on graph **B**: #, *P* < 0.001 for Δ*sgl1*/isotype → WT vs PBS/isotype → WT or Δ*sgl1*/anti-CD4 → WT vs. PBS/anti-CD4 → WT.

### Therapeutic administration of HK C. neoformans Δsgl1 post WT challenge reduces WT cells in the lungs of vaccinated mice

From our previous studies and present work, complete host protection in either immunocompetent or immunocompromised *C. neoformans* Δ*sgl1*-vaccinated mice has always been associated with persistent WT fungal cells remaining in the lungs post WT challenge [52] (T.G. Normile, T.H. Chu, B.S. Sheridan, and M. Del Poeta, submitted for publication). In those studies, we showed that the lung fungal burden at days 45, 75, and 105 post WT challenge was nearly identical at all timepoints, and histopathology at these timepoints displayed a decreased percentage of inflamed lung tissue and increased formation isolated nodules of contained yeast cells [52]. In this study, we found for the first time that vaccination with two subsequent doses of HK *C. neoformans* Δ*sgl1* results in a significant decrease of lung fungal burden compared to live *C. neoformans* Δ*sgl1*-vaccination for both immunocompetent and CD4-deficient mice (**Fig. 1D**).

Moreover, vaccinated mice that received more than one WT challenge displayed a significant reduction in the lung fungal burden compared to mice that were received only one WT challenge (**Fig. 3C**). Because subsequent WT challenges decreased the lung fungal burden, we asked if administration of our vaccine could be used as a therapeutic strategy and administered after the WT challenge.

To investigate the therapeutic potential of HK *C. neoformans* Δ*sgl1* administration in *C. neoformans* Δ*sgl1*-vaccinated mice, immunocompetent and CD4-deficient mice were challenged with the WT strain first and then received either 1 or 2 subsequent administrations of HK *C. neoformans* Δ*sgl1* (see experimental design schematic: **Fig. 4A**). We found a significant decrease in the lung fungal burden after therapeutic administration of HK *C. neoformans* Δ*sgl1* in both immunocompetent and CD4-deficient mice (**Fig. 4B**). From the baseline lung fungal burden on day 30 post challenge, there was a significantly greater lung burden in CD4-deficient mice compared to the isotype-treated as we have seen previously. In addition, the lung fungal burden in mice that were treated with either 1 or 2 administrations of PBS (control groups) on days 30 and 45, respectively, was nearly identical to the baseline lung fungal burden (**Fig. 4B**). Interestingly, there was a significant decrease in the lung burden in mice that received 1 or 2 administrations of HK *C. neoformans* Δ*sgl1* post WT challenge compared to the PBS-treated groups at those timepoints as well as in the lung burdens between mice that received 1 or 2 administrations of HK *C. neoformans* Δ*sgl1* (**Fig. 4B**). Together, these data suggest that therapeutic administration of HK *C. neoformans* Δ*sgl1* post WT challenge significantly reduces the number of persistent WT yeast in the lungs of vaccinated mice.

**Figure 5.**
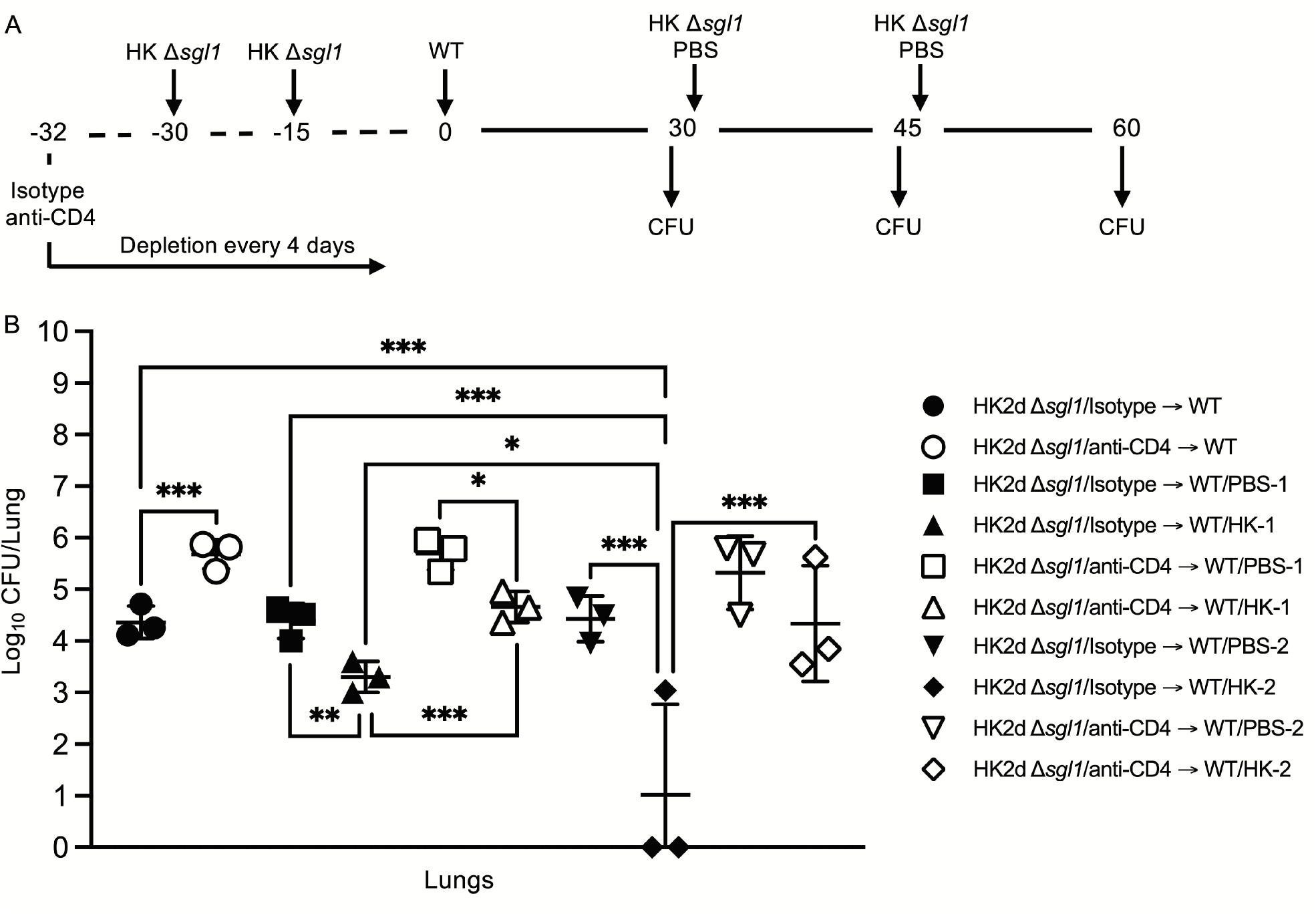
Administration of heat-killed (HK) *C. neoformans* Δ*sgl1* post wild-type (WT) challenge significantly reduces persistent WT yeast from the lungs of vaccinated mice. **A**. Experimental design schematic. CBA/J mice (n=9 mice/group) were administered either isotype or anti-CD4 antibody prior to vaccination with two identical doses of 5x10^7^ HK Δ*sgl1* on days - 30 and -15, and all mice were challenged with 5x10^5^ *C. neoformans* WT on day 0. After 30 days post WT challenge, 3 mice/group (HK2d Δ*sgl1*/isotype → WT and HK2d Δ*sgl1*/anti-CD4 → WT) were sacrificed for lung fungal burden determination, and the remainder of the mice were then administered either 5x10^7^ HK Δ*sgl1* or PBS. Fifteen days later on day 45, 3 mice/group were sacrificed for lung fungal burden determination (HK2d Δ*sgl1*/isotype → WT/PBS-1, HK2d Δ*sgl1*/isotype → WT/HK-1, HK2d Δ*sgl1*/anti-CD4 → WT/PBS-1, and HK2d Δ*sgl1*/anti-CD4 → WT/HK-1), and the remainder of the mice were then administered either 5x10^7^ HK Δ*sgl1* or PBS. Fifteen days later on day 60, 3 mice/group were sacrificed for lung fungal burden determination (HK2d Δ*sgl1*/isotype → WT/PBS-2, HK2d Δ*sgl1*/isotype → WT/HK-2, HK2d Δ*sgl1*/anti-CD4 → WT/PBS-2, and HK2d Δ*sgl1*/anti-CD4 → WT/HK-2). **B.** Endpoint organ fungal burden was quantified in the lungs for mice sacrificed on days 30, 45, and 60 (n=3 mice/group/timepoint). Graphed data represent the mean +/- SD (**B**). Significance was determined by an Ordinary one-way ANOVA using Tukey’s multiple comparisons test for *P* value adjustment and is denoted as *, *P* < 0.05; **, P < 0.01; ***, *P* < 0.005.

### Therapeutic administration of live or HK C. neoformans Δsgl1 post WT challenge significantly prolongs survival in unvaccinated mice

Because we observed the efficacious therapeutic potential of HK *C. neoformans* Δ*sgl1* administration post WT challenge in vaccinated mice, we then asked if therapeutic administration of HK *C. neoformans* Δ*sgl1* post WT challenge was useful in naïve, unvaccinated mice.

We tested this hypothesis by challenging naive mice with the WT strain and then administered HK *C. neoformans* Δ*sgl1*, live *C. neoformans* Δ*sgl1*, or PBS on either day 3 or day 7 and monitored for survival (experimental design schematic: **Fig. 5A**). While all mice administered PBS fatally succumbed to infection, all mice administered HK *C. neoformans* Δ*sgl1* or live *C. neoformans* Δ*sgl1* on day 3 post WT challenge survived to the experimental endpoint (**Fig. 5B**). In addition, mice administered HK *C. neoformans* Δ*sgl1* or live *C. neoformans* Δ*sgl1* on day 7 post WT challenge exhibited a 70% and 60% survival rate at the experimental endpoint, respectively. Nevertheless, there were no differences in the lung fungal burden between any of the surviving groups (**Fig. 5B**). Of note, all surviving mice displayed extrapulmonary dissemination of the WT strain to the brain. Interestingly, mice that received therapeutic administration of live or HK *C. neoformans* Δ*sgl1* on day 3 had fewer brain CFU compared to mice administered on day 7 (**Fig. 5C**). Overall, these data suggest that live or HK *C. neoformans* Δ*sgl1* aids to significantly prolong the survival of mice from fatal WT infection.

**Figure 6.**
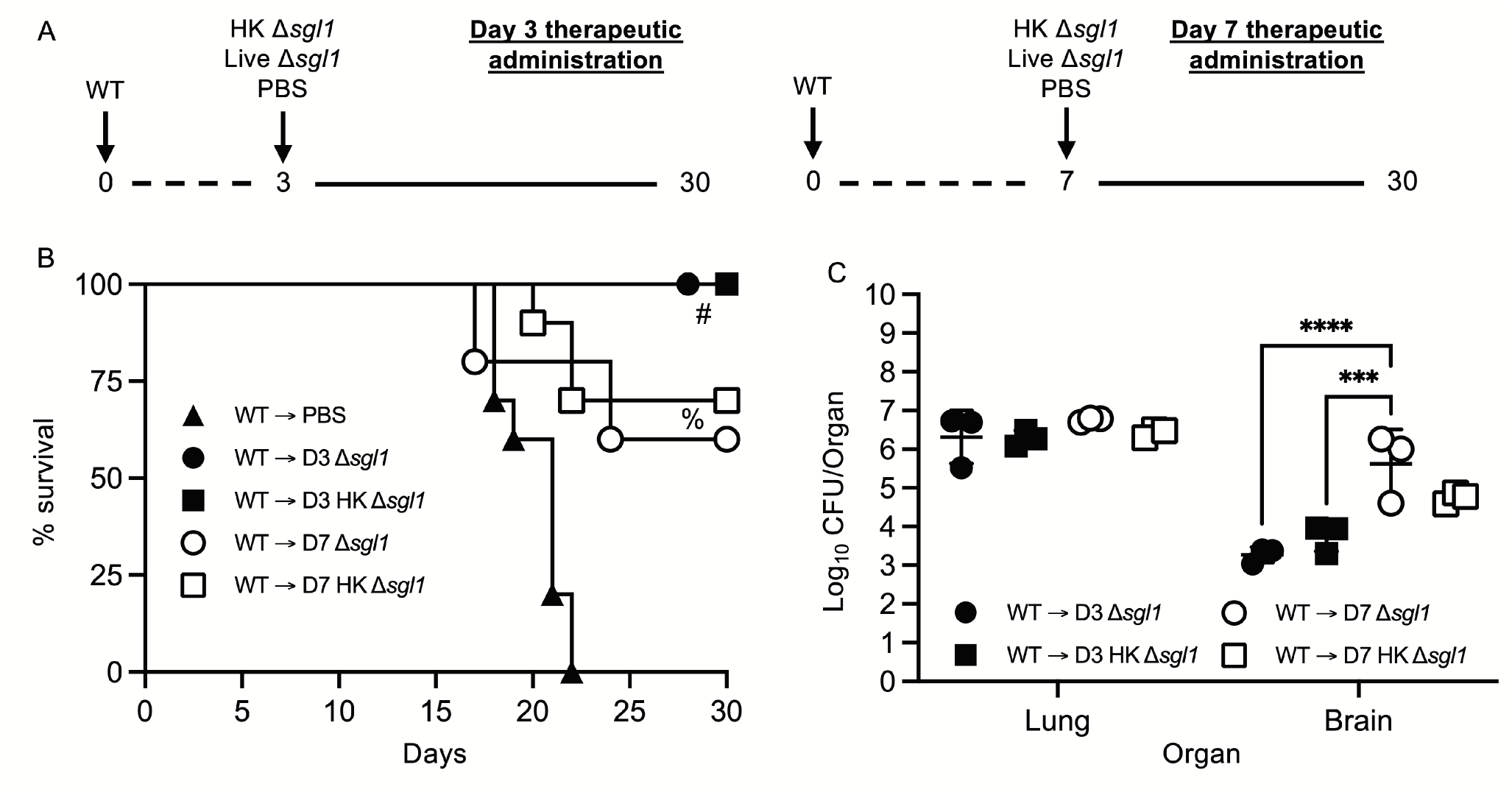
Therapeutic administration of live or heat-killed (HK) *C. neoformans* Δ*sgl1* significantly prolongs host survival to lethal WT infection. **A**. Experimental design schematic. CBA/J mice (n=10 mice/group) were challenged with 1x10^5^ *C. neoformans* WT on day 0. WT challenged mice were administered 5x10^7^ HK Δ*sgl1*, 5x10^5^ Live Δ*sgl1*, or PBS on either day 3 (D3) or day 7 (D7) and monitored for survival until the experimental endpoint. **C.** Endpoint organ fungal burden was assessed in the lungs and brains of surviving mice (n=3 mice/group). Graphed data represent the survival percentage of WT challenged mice (**B**) or the mean +/- SD (**C**). Significance was determined by a two-way ANOVA using Šídák’s multiple comparisons test for *P* value adjustment (**C**) and significance is denoted as ***, *P* < 0.005; ****, *P* < 0.001. Survival significance was determined by the Mantel-Cox log-rank test (**B**) and denoted on graph **B**: #, *P* < 0.001 for WT → D3 Δ*sgl1* or WT → D3 HK Δ*sgl1* vs. WT → PBS; %, *P* < 0.001 for WT → D7 Δ*sgl1* or WT → D7 HK Δ*sgl1* vs. WT → PBS.

### Vaccination with live or HK C. neoformans Δsgl1 protects mice from fatal infection by reactivation of latent cryptococcosis via immunosuppression

We have now shown that live or HK *C. neoformans* Δ*sgl1* can be effectively used both preventatively (**Figs. 2-4**) and therapeutically (**Figs. 5-6**) to elicit robust host protection. However, *C. neoformans* is not only a primary pathogen since fungal cells can be contained within lung granulomas in immunocompromised hosts for extensive periods of time but immunosuppressive conditions, such as CD4-lymphopenic HIV/AIDS patients, can cause granuloma breakdown, latent cell proliferation, and brain dissemination potentially resulting in fatal meningoencephalitis [11, 31]. Because of this, we investigated the ability of *C. neoformans* Δ*sgl1* to protect mice from cryptococcal reactivation from a lung granuloma.

To test this, mice were intranasally inoculated with *C. neoformans* Δ*gcs1*, a mutant strain lacking glucosylceramide synthase, that has been previously reported to induce pulmonary granuloma formation in mice over 30 days. At 30 days post Δ*gcs1* administration, we administered live *C. neoformans* Δ*sgl1*, HK *C. neoformans* Δ*sgl1*, or PBS. After another 30 days, all groups of mice underwent either corticosteroid-induced immunosuppression to induce leukopenia or CD4^+^ T cell depletion to induce CD4 lymphopenia, and mice were monitored for survival (simplified experimental design schematic: **Fig. 6A**; detailed experimental design schematic: **Supplementary Fig. S1**). Extraordinarily, we observed that mice administered live *C. neoformans* Δ*sgl1* or HK *C. neoformans* Δ*sgl1* exhibited a 75% and 62.5% survival rate, respectively, at the experimental endpoint post corticosteroid-induced immunosuppression, while all PBS-treated mice fully succumbed to fatal reactivation (**Fig. 6B**). Similarly, mice administered live *C. neoformans* Δ*sgl1* or HK *C. neoformans* Δ*sgl1* exhibited a 100% and 87.5% survival rate, respectively, at the experimental endpoint post CD4^+^ T cell depletion, which were significantly greater than the PBS-treated mice that displayed a 37.5% survival rate (**Fig. 6C**). These data suggest that vaccination with live or HK *C. neoformans* Δ*sgl1* can be used to protect the host from cryptococcal reactivation from a lung granuloma in the event that they become immunocompromised.

**Figure 7.**
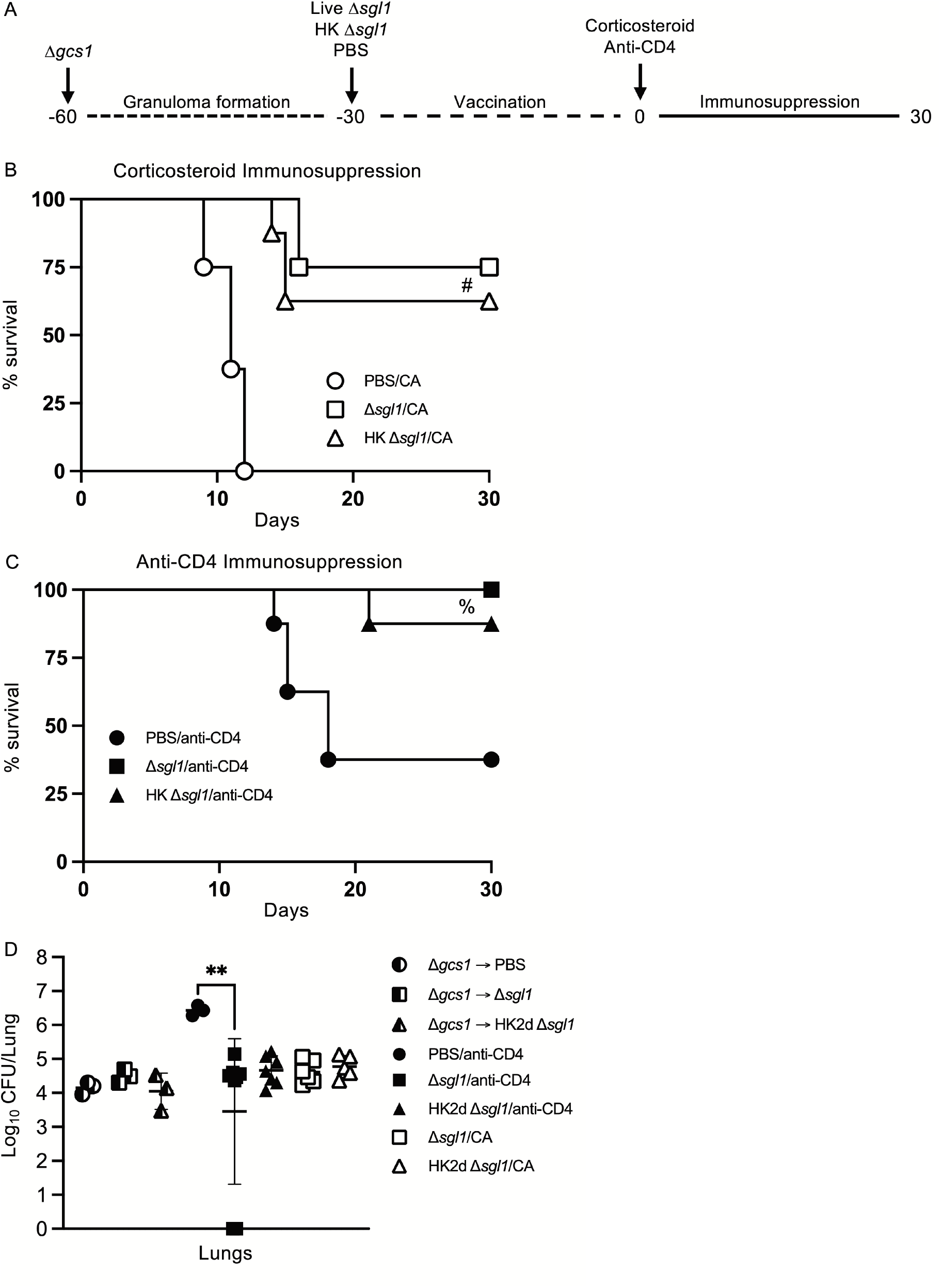
Vaccination with either live or heat-killed (HK) *C. neoformans* Δ*sgl1* protects mice from lethal reactivation infection post immunosuppression. **A.** Experimental design schematic (a more detailed schematic can be found in Supplementary Figure S1). B-C. CBA/J mice were infected with 5x10^5^ *C. neoformans* Δ*gcs1* on day -60 to induce lung granuloma formation. After 30 days, mice were administered either 5x10^5^ Live Δ*sgl1* or PBS on day -30 or 5x10^7^ HK Δ*sgl1* on days -30 and -15. Finally on day 0, all groups of mice underwent continuous immunosuppressive treatment with either the corticosteroid cortisone acetate (CA) (B) or anti-CD4 antibody (C) to cause reactivation of the latent *C. neoformans* Δ*gcs1* yeast contained within the lung granulomas and assessed for survival over 30 days. D. Endpoint lung fungal burden comparison in mice pre-immunosuppression on day 0 (Δ*gcs1* → PBS and Δ*gcs1* → Δ*sgl1*) (n=3 mice/group) and post-immunization on day 30 for CA-treated mice (Δ*sgl1*/CA and HK2d Δ*sgl1*/CA) (n=6-7 mice/group) and anti-CD4-treated mice (PBS/anti-CD4, Δ*sgl1*/anti-CD4, and HK2d Δ*sgl1*/anti-CD4) (n=9-10 mice/group). Graphed data represent the survival percentage of mice (B, C) and the mean +/- SD (D). Significance was determined by an Ordinary one-way ANOVA using Tukey’s multiple comparisons test for *P* value adjustment (D) and is denoted as **, *P* < 0.01. The Mantel-Cox log-rank test was used to determine survival significance (B, C) and denoted on each graph: B: #, *P* < 0.001 for either Δ*sgl1*/CA or HK Δ*sgl1*/CA vs. PBS/CA; C: %, *P* < 0.01 for Δ*sgl1*/anti-CD4 or HK Δ*sgl1*/anti-CD4 vs. PBS/anti-CD4.

To examine the efficacy of our vaccine strategy in the experimental reactivation model, the endpoint lung fungal burden in mice pre-immunosuppression (day 0) was compared to the fungal burden in the lungs of mice that survived until the experimental endpoint post-immunosuppression (day 30). We first observed there were no differences in the lung fungal burdens between any of the groups pre-immunosuppression (**Fig. 6D**). Interestingly, there were no statistical differences between the endpoint lung fungal burdens post-immunosuppression in *neoformans* Δ*sgl1*-vaccinated mice or between the endpoint lung fungal burdens in mice pre-immunosuppression compared to in mice post-immunosuppression. In fact, the only observed statistically significant difference was between the surviving PBS-treated CD4-deficient mice and live *C. neoformans* Δ*sgl1*-vaccinated mice, which further supports that vaccination with *C. neoformans* Δ*sgl1* protects mice from lethal reactivation upon immunosuppression (**Fig. 6D**). Comparably, all surviving PBS-treated mice displayed significantly greater brain dissemination compared to vaccinated mice, which were almost fully absent of any brain fungal burden (**Supplementary Fig. S2**). Overall, these data suggest that administration of live or HK *C. neoformans* Δ*sgl1* inhibits the proliferation of dormant fungal cell in the lung granuloma and protects from extrapulmonary dissemination upon immunosuppression.

## Discussion

In the current study, we have presented ample evidence on the highly efficacious use of HK *C. neoformans* Δ*sgl1* in conferring robust host protection in three separate models of vaccination against cryptococcosis during immunosuppression. More specifically, we have shown that: i) preventative vaccination with 2 doses of HK *C. neoformans* Δ*sgl1* conferred complete host protection to lethal challenge with decreased endpoint lung fungal burden compared to live cell vaccination; ii) therapeutic administration of HK *C. neoformans* Δ*sgl1* post WT challenge resulted in a continual decrease in the lung fungal burden with each subsequent vaccine administration, conferring significantly increased survival rate; and iii) our vaccination strategy prevented cryptococcal reactivation from a lung granuloma, by inhibiting proliferation of latent fungal cells and improving survival upon immunosuppression.

Host protection was both concentration- and dose-dependent requiring 2 subsequent administrations of 5x10^7^ HK *C. neoformans* Δ*sgl1*. The requirement for multiple doses has been seen with other HK vaccine-inducing mutant strains [55, 56], while others required only one dose [58, 59]. Because all these studies including our current work use a similar vaccine concentration between 1x10^7^ and 5x10^7^, the only difference for the single dose requirement was the use of the KN99 WT strain compared to the H99 WT strain. Nevertheless, all studies on HK vaccine-inducing mutants, including this current work, report 100% protection to the lethal WT challenge. However, the true standout characteristic for a clinically relevant vaccine formulation is the ability to induce protection in a model most associated with a disease, which is CD4- deficiency for cryptococcosis [36, 40].

In comparison to our present findings where we report 100% protection in CD4- deficiency with HK *C. neoformans* Δ*sgl1* vaccination, the only other HK vaccine-inducing mutant to demonstrate protection during CD4-deficiency was from Wang and colleagues using a HK F-box protein (Δf*bp1*) mutant strain [55]. Similarly, both HK *C. neoformans* Δ*sgl1* and the HK Δf*bp1* mutants demonstrated complete protection in both immunocompetent and CD4- deficient CBA/J mice, although differences between the two are noteworthy. First, our present work with HK *C. neoformans* Δ*sgl1* resulted in a ∼1 log lower endpoint lung fungal burden for isotype-treated mice compared to isotype-treated mice in the report by Wang and colleagues (although the endpoint lung fungal burden for CD4-deficient mice was nearly identical) [55] (**Fig. 1**). Second, we observed a complete lack of any extrapulmonary dissemination in mice vaccinated with HK *C. neoformans* Δ*sgl1*, while several mice displayed fungal CFU in the brain and spleen in the study by Wang and colleagues [55]. Finally, the WT challenge dose used in our work was 15x greater than used by Wang and colleagues.

With regards to our vaccine, we aimed to test the rigor and robustness of *C. neoformans* Δ*sgl1* in the preventative model of vaccination via functional alterations to our experimental design. Since T cell mediated immunity is a well-established keystone of anti-cryptococcal immunity [60, 61] as well as the need for either CD4^+^ or CD8^+^ T cells for *C. neoformans* Δ*sgl1* protection [52], these alterations focused upon memory T cells. The first alteration involved a 3- fold increase in the time between vaccination and WT challenge, where vaccination began 90 days prior to WT challenge for both live and HK *C. neoformans* Δ*sgl1*. All immunocompetent and CD4-deficient mice vaccinated with live *C. neoformans* Δ*sgl1* survived the lethal WT challenge, and a respective 90% and 70% survival was observed in mice vaccinated with HK *C. neoformans* Δ*sgl1* (**Fig. 2A, C**). The protection observed in the extended rest period suggests the induction of long-lived memory T cells post vaccination with *C. neoformans* Δ*sgl1*. Future immunophenotyping assays will be aimed to define the type of circulating memory T cells, such as central memory, tissue-resident memory, or effector memory.

Complete host protection was not observed in 100% of the CD4^+^ deficient mice when they were vaccinated with HK *C. neoformans* Δ*sgl1* 90 days prior to WT challenge. This suggests that the immunological memory induced was either less robust or shorter-lived compared to vaccination with the live cell strain. It is noteworthy to mention that the WT challenge dose was doubled in this experimental design due to the increased age of the mice at the time of challenge. However, the decreased length of antigen encounter using HK mutant strains may have potentially resulted in less robust naïve T cell stimulation and fewer memory T cells following the contraction phase [54, 62]. Optimization of the dosing regimen will be required in future studies. Potential adjustments could include increasing the number of doses, increasing the time between the first and second dose, or altering the concentrations to induce more robust immunity with a lower first dose and a greater second dose.

The second functional alteration to the preventative vaccination model experimental design was to increase the number of WT challenges administered to vaccinated mice. During chronic infections, such as when fungal cells are persisting in the lungs, T cells may become tolerized to antigens remaining alive for extended periods of time in a hyporesponsive state known as T cell anergy [57, 63]. Mice vaccinated with *C. neoformans* Δ*sgl1* exhibited the opposite, however. First, all mice that received two or three subsequent WT challenges exhibited 100% survival even during CD4^+^ T cell deficiency (**Fig. 3B**). Second, the endpoint lung fungal burden in mice that received at least 2 WT challenges displayed a ∼2 log decrease compared to mice that received only 1 WT challenge (**Fig. 3C**). This suggests the efficacy observed with the functional alterations in the preventative model during vaccination with *C. neoformans* Δ*sgl1* elicits long-lived, non-exhaustive T cell memory with increasing clonal functionality upon subsequent WT encounters. Future work will address a phenotypic and functional characterization comparing T cells from mice administered one WT challenge with mice administered more than one challenge.

Persistent fungal cells remaining in the lungs post WT challenge in *C. neoformans* Δ*sgl1*- vaccinated mice have been an observable facet in all experimental variations in this study and previous work from our lab. Moreover, fungal cell persistence post WT challenge in vaccinated mice has been reported in other cryptococcal vaccine studies as well [55, 56, 58, 59, 64, 65]. Because we observed a decrease in the lung fungal burden after a second WT challenge (**Fig. 3C**), we investigated the immunotherapeutic ability of HK *C. neoformans* Δ*sgl1* to decrease further WT fungal cells remaining in the lungs. The first administration of HK *C. neoformans* Δ*sgl1* significantly decreased the persistent fungal burden to a similar degree as mice that received a second WT challenge (**Fig. 4B** and **Fig. 3C**). Interestingly, mice that received a second administration of HK *C. neoformans* Δ*sgl1* significantly decreased the remaining fungal cells to an even further extent compared to mice that were administered PBS or mice that received only 1 therapeutic dose of HK *C. neoformans* Δ*sgl1* (**Fig. 4B**). In fact, 2 of the 3 mice fully cleared the WT fungal cells from the lungs. Thus, HK *C. neoformans* Δ*sgl1* exhibits robust immunotherapeutic potential in previously vaccinated immunocompetent mice.

Collectively, the therapeutic potential of HK *C. neoformans* Δ*sgl1* administration has demonstrated highly efficacious host protection in both previously vaccinated (**Fig. 4**) and unvaccinated mice (**Fig. 5**). While this adds an entirely new dimension to our vaccine, immunotherapeutic administration is scarce in the literature with only a few other reports. The first immunotherapeutic study utilized P13, an antigenic peptide mimotope of the cryptococcal capsular GXM conjugated to either tetanus toxoid or diphtheria toxoid [66, 67]. Immunization with P13 after an otherwise lethal challenge significantly prolonged survival, yet all mice soon succumbed to fatal infection [66]. Similarly, Datta and colleagues established a model of chronic infection in mice and administration of P13 significantly prolonged host survival compared to control mice, but again all mice soon succumbed to fatal infection [67]. In addition to the P13 conjugate vaccine, a TNFα-expressing adenoviral vector was also utilized post lethal WT challenge [68]. Although survival was not assessed, the authors reported a significant decrease in lung fungal burden, increased IFNγ levels, and a significant increase in macrophage and neutrophil recruitment to the lungs. Overall, in addition to the robust efficacy in the preventative model of vaccination, *C. neoformans* Δ*sgl1* has now been shown to possess unrivaled immunotherapeutic potential adding to the clinical significance of our vaccine.

Although both the prevention and therapeutic models increase the novelty and translational potential of our vaccine, we have demonstrated vaccine-induced host protection against lethal infection due to reactivation of latent fungal cells upon immunosuppressive treatments (**Fig. 6B, C**). To our knowledge, this is the first time a vaccine against the reactivation infection has been reported in the literature. Previous work in our lab had shown that mice treated with FTY720, a prescribed treatment for relapsing remitting multiple sclerosis, was linked to granuloma breakdown with a disorganization of the peripheral macrophages with a shift towards an M2 polarized state [11]. In addition, our findings also validate the reactivation model, as it showed that the *C. neoformans* Δ*gcs1*-induced granuloma in mice can lose integrity upon immunosuppression resulting in fungal proliferation in the lung, brain dissemination, and ultimately death.

In fact, clinical cases can occur due to the reactivation of granuloma-contained fungal cells from either immunosuppression or comorbidities (HIV/AIDS progression) [27, 69]. Because of this, we tested our vaccination strategy in this mouse model during prolonged corticosteroid-induced immunosuppression as well as CD4-deficiency. We observed a 70% and 60% survival rate in mice vaccinated with live or HK *C. neoformans* Δ*sgl1*, respectively, at the endpoint after corticosteroid-induced immunosuppression with cortisone acetate (**Fig. 6B**), and a 100% and 90% survival rate in mice vaccinated with live or HK *C. neoformans* Δ*sgl1* at the endpoint after depletion of CD4^+^ T cells (**Fig. 6C**). Interestingly, the corticosteroid-induced immunosuppression was more lethal than the depletion of CD4^+^ T cells, which may be attributed to the mechanism of immunosuppression. Corticosteroid-induced immunosuppression induces leukopenia, inhibits phagocytosis, and decreases antigen presentation capabilities [70, 71], while depletion of CD4^+^ T cells ablates circulating CD4^+^ lymphocytes. So, we speculate that the difference in lethality of the infection may be the speed at which the immunosuppression took effect.

Although there was an observed difference in survival between the two modes of immunosuppression, the endpoint lung fungal burden between the two modes of immunosuppression were nearly identical (**Fig. 6D**). In fact, there were also no differences between the endpoint lung fungal burden of mice pre-immunosuppression and *C. neoformans* Δ*sgl1*-vaccinated mice post-immunosuppression. This suggests that vaccination with either live or HK *C. neoformans* Δ*sgl1* controls the proliferation of the latent fungal cells in the lungs even after the immunosuppressive regime. This is further supported from the endpoint lung fungal burden in unvaccinated CD4-deficient mice being significantly greater than the lung fungal burden in the vaccinated mice, which indicate that fungal cells extensively proliferate in unvaccinated mice upon immunosuppression. The same was observed for extrapulmonary dissemination to the brain (**Supplementary Fig. S2**). While there were only 1-2 *C. neoformans* Δ*sgl1*-vaccinated mice that displayed fungal dissemination, all the surviving unvaccinated mice had significant fungal burden in the brain. Overall, vaccination with either live or HK *C. neoformans* Δ*sgl1* demonstrated remarkable efficacy in this cryptococcal model of reactivation.

In conclusion, we have shown here that HK *C. neoformans* Δ*sgl1* demonstrates a highly efficacious vaccine candidate that goes beyond the canonical preventative model of primary disease prevention. We have expanded not only to a more clinically relevant HK formulation but also to additional models of vaccine strategies to protect against cryptococcosis during CD4- deficiency, including using our vaccine as a therapeutic mean and using our vaccine to prevent reactivation of a latent infection upon immunodepression. Here forth, the tools for investigation into the protective immunity against fungal reactivation from pulmonary granulomas in mice are now available, which greatly opens future possibilities to significantly add to this completely absent portion of the literature.

## Materials and Methods

### Fungal strains and heat killed (HK) yeast

The fungal strains used in this study were wild-type (WT) *C. neoformans* var. *grubii* strain H99, *C. neoformans* Δ*sgl1*, a mutant strain accumulating sterylglucosides developed by our group [45], and *C. neoformans* Δ*gcs1*, a mutant strain lacking glucosylceramide synthase [72]. For all experiments, fungal strains were freshly recovered from a -80°C freezer stock on YPD agar at 30°C for 3-4 days. An isolated colony was added to 10ml of YPD broth and grown for 16-18hr with shaking, washed three times in sterile PBS, counted with a hemocytometer, and resuspended in sterile PBS at the desired concentration. For HK strains, the desired concentration of live yeast was resuspended in PBS and added to an 80°C heat block for 1hr. All HK strains were confirmed to be fully dead by plating the mixture on YPD plates at 30°C for 4 days and observing no growth.

### Mice and ethical statement

Both male and female CBA/J mice were purchased from Envigo. All animals were housed under specific pathogen free conditions and had access to food and water *ad libitum*. Mice were allowed one week to acclimate upon arrival before any procedures began. All animal procedures were approved by the Stony Brook University Institutional Animal Care and Use Committee (protocol no. 341588) and followed the guidelines of American Veterinary Medical Association.

### In vivo infections and organ fungal burden

All primary infections and immunizations were carried out in both male and female CBA/J mice 4-6 weeks old. Mice were first intraperitoneally (IP) anesthetized with a ketamine/xylazine solution (95mg of ketamine and 5mg of xylazine per kg of animal body weight). Anesthetized mice were then intranasally (IN) injected with the desired concentration of the specified yeast cells in 20μl of PBS. For fungal burden analysis, mice were euthanized via CO_2_ inhalation on pre-determined timepoints. The lungs, brain, spleen, kidneys, and liver were removed, homogenized in 10ml of sterile PBS using a Stomacher 80 blender (Seward, UK), and serial dilutions were grown on YPD plates at 30°C for 3-4 days before being counted and total organ burden calculated.

### Immunosuppression treatments

Cortisone 21-acetate (CA) (Sigma; cat # C3130) was used to induce leukopenia. Mice were sub-cutaneously administered 250mg/kg/mouse CA in PBS every other day for a set timeline. IP administration of anti-CD4 monoclonal antibody (clone: GK1.5; BioXCell) was used to deplete mice of CD4^+^ T cells. Antibody dilutions were prepared from the stock solution in PBS each time. Mice were administered 400μg/100μl every 4 days for the duration of the experiment to maintain cell depletion. Control group mice were administered isotype-matched antibody at the same concentration and administration timeline.

### Vaccination strategies and survival studies

Three different vaccination models were used in this study. For survival studies, any animal that appeared to be moribund, exhibited labored breathing or neurological infection, or had lost more than 20% body weight was euthanized via CO_2_.

(1) For the prevention model, mice were IN injected with 5x10^5^ live *C. neoformans* Δ*sgl1* in 20μl of PBS, 5x10^5^, 5x10^6^, or 5x10^7^ HK *C. neoformans* Δ*sgl1* in 20μl of PBS, or 20μl of sterile PBS (unvaccinated controls) 30 days prior to WT challenge unless stated otherwise in the figure caption. Mice were challenged with 5x10^5^ *C. neoformans* WT in 20μl of PBS unless stated otherwise in the figure caption and monitored daily until the pre-determined experimental endpoint.
(2) For the therapeutic model, live or HK *C. neoformans* Δ*sgl1* was used to treat vaccinated or unvaccinated mice post WT challenge. In vaccinated mice, immunocompetent or CD4-deficient mice were administered two subsequent doses of 5x10^7^ HK *C. neoformans* Δ*sgl1* on days -30 and -15, and mice were challenged with the WT strain on day 0. Mice were administered additional doses of 5x10^7^ HK *C. neoformans* Δ*sgl1* on days 30 and 45 to reduce WT cells that persist in the lungs of vaccinated mice. In unvaccinated mice, mice were first challenged with 1x10^5^ *C. neoformans* WT strain on day 0. WT challenged mice were treated with 5x10^5^ live *C. neoformans* Δ*sgl1* in 20μl of PBS, 5x10^7^ HK *C. neoformans* Δ*sgl1* in 20μl of PBS, or 20μl of sterile PBS (controls) on either day 3 or day 7 post challenge and assessed for survival until day 30.
(3) For the reactivation model, we assessed whether vaccination with *C. neoformans* Δ*sgl1* could protect mice from lethal reactivation infection from latent fungal cells. First, mice were IN injected with 5x10^5^ *C. neoformans* Δ*gcs1*, an avirulent mutant that has been shown to induce lung granuloma formation that recapitulates a human pulmonary granuloma [72], on day - 60. After 30 days, mice were IN injected with 5x10^5^ live *C. neoformans* Δ*sgl1*, 5x10^7^ HK *C. neoformans* Δ*sgl1*, or 20μl of sterile PBS (unvaccinated controls) on day -30 (and a second dose of 5x10^7^ HK *C. neoformans* Δ*sgl1* on day -15). To induce reactivation of latent *C. neoformans* Δ*gcs1*, mice were immunosuppressed via administration of either corticosteroids (cortisone 21- acetate) or depleted of CD4^+^ T cells via IP injection of a monoclonal antibody at set timelines beginning on day 0. Mice were monitored for survival over 30 days.

### Statistical analysis

All analyses were performed using GraphPad Prism 9 software. The sample size, statistical analysis, and statistical significance is described in each figure caption. The Mantel-Cox log-rank test was used to calculate significance for survival studies. A two-tailed unpaired *t* test was used to calculate statistical significance between two samples, and either an ordinary one-way ANOVA using Tukey’s multiple comparisons test for *P* value adjustment or a two-way ANOVA using Šídák’s multiple comparisons test for *P* value adjustment was used to calculate statistical significance between more than two samples.

## Acknowledgments

This work was supported by the National Institute of Health (NIH) grants AI136934 (MDP), AI116420 (MDP), and AI125770 (MDP), and by a Merit Review Grant I01BX002924 (MDP) from the Veterans Affairs (VA) Program. Maurizio Del Poeta is a recipient of the Research Career Scientist (RCS) Award (IK6 BX005386) and a Burroughs Welcome Investigator in Infectious Diseases.

## Author contributions

TGN and MDP took part in the conceptualization of this study as well as the writing and finalization of the manuscript. TGN performed all animal infections, experimental procedures, statistical analysis, and figure generation.

